# Ectopic expression of the germline transcription factor LSL-1 contributes to developmental delay following failed maternal epigenetic reprogramming

**DOI:** 10.64898/2026.08.28.747910

**Authors:** Benjamin Nguyen, Zaynab Massenburg, Josh Labus, Sundas Johnson, Sarah D. Blancher, Miyah Jones, Brandon S. Carpenter

## Abstract

Proper transmission of cell identity between generations requires maternal epigenetic reprogramming mechanisms that prevent inappropriate inheritance of lineage-specific transcriptional programs. In *Caenorhabditis elegans*, loss of the H3K4me1/2 demethylase SPR-5 and the H3K9 methyltransferase MET-2 results in ectopic expression of germline genes in somatic tissues and severe developmental delay. Previous studies demonstrated that the chromatin regulator MES-4 contributes to these defects, but whether germline-specific transcription factors also participate in the ectopic transcriptional program remained unclear. Here, we investigated the role of the germline transcription factor LSL-1 in animals lacking SPR-5 and MET-2. We found that genes normally regulated by LSL-1 in the germline are significantly overrepresented among genes ectopically expressed in the soma of *spr-5; met-2* progeny. Consistent with this observation, an endogenously tagged LSL-1 protein became ectopically expressed throughout somatic tissues when maternal SPR-5 and MET-2 activity was disrupted. Furthermore, depletion of LSL-1 partially suppressed the developmental delay observed in *spr-5; met-2* mutants. Transcriptomic analyses revealed extensive overlap between MES-4- and LSL-1-dependent transcriptional programs, with most LSL-1-dependent genes also requiring MES-4. Notably, ectopic expression of *lsl-1* itself depended on MES-4, suggesting that LSL-1 functions downstream of MES-4. Genes dependent on LSL-1 were strongly enriched for germline-associated expression programs and included previously identified direct LSL-1 targets. Together, our findings support a model in which MES-4 promotes ectopic expression of LSL-1, which in turn contributes to a shared germline-associated transcriptional program and developmental delay following failed maternal epigenetic reprogramming. These results demonstrate how lineage-restricted transcription factors cooperate with inherited chromatin states to reinforce aberrant transcriptional programs and disrupt cell fate boundaries.

## INTRODUCTION

Cell fate determination during embryonic development requires the coordinated establishment and maintenance of lineage-specific gene expression programs. Epigenetic mechanisms, including histone methylation, play central roles in restricting transcriptional programs to appropriate cell types and preventing the propagation of inappropriate developmental states (Burton and Torres-Padilla 2014; Frum and Ralston 2015). In *Caenorhabditis elegans*, maternally supplied chromatin regulators reset transcriptional memory at fertilization to ensure proper separation of germline and somatic identities across generations (Katz et al. 2009; Greer et al. 2014; Kerr et al. 2014). Failure of this maternal reprogramming process results in the persistence of inherited chromatin states and the inappropriate activation of germline-specific genes in somatic tissues, leading to developmental defects (Greer et al. 2014; Kerr et al. 2014; Carpenter et al. 2021).

Maternal reprogramming in *C. elegans* is mediated in part by the H3K4me1/2 demethylase SPR-5 and the H3K9me1/2 methyltransferase MET-2. Together, these enzymes remove activating chromatin marks and establish transcriptionally repressive chromatin states during early embryogenesis. Loss of both SPR-5 and MET-2 results in synergistic accumulation of inherited H3K4 methylation and widespread ectopic expression of germline genes in somatic tissues, ultimately causing severe developmental delay (Katz et al. 2009; Greer et al. 2014; Kerr et al. 2014; Carpenter et al. 2021). Previous work demonstrated that this phenotype depends in part on the H3K36 methyltransferase MES-4, which maintains transcriptional memory of germline-expressed genes across generations and contributes to their inappropriate activation in the soma when maternal reprogramming fails (Furuhashi et al. 2010; Rechtsteiner et al. 2010; Carpenter et al. 2021).

While inherited chromatin states can create permissive conditions for aberrant gene expression, sustained activation of lineage-specific transcriptional programs often requires sequence-specific transcription factors. In many developmental contexts, transcription factors cooperate with chromatin-based regulatory mechanisms to establish and reinforce stable patterns of gene expression that maintain cell identity (Graf and Enver 2009; Iwafuchi-Doi and Zaret 2014). We therefore considered the possibility that germline-specific transcription factors may help promote ectopic germline gene expression in the soma of *spr-5; met-2* mutants once aberrant chromatin states have been established. Recently, Rodríguez-Crespo and colleagues identified LSL-1 as a major activator of germline transcription that is required for normal germ cell development and expression of numerous germline genes (Rodriguez-Crespo et al. 2022). LSL-1 directly binds and activates genes involved in meiosis, chromosome pairing, and germ cell identity, establishing it as a central regulator of germline transcriptional programs (Rodriguez-Crespo et al. 2022). Given the widespread ectopic expression of germline genes in *spr-5; met-2* mutant soma, we hypothesized that inappropriate expression of LSL-1 may contribute to ectopic germline transcription when maternal reprogramming is disrupted.

Here, we show that LSL-1 is ectopically expressed in the soma of *spr-5; met-2* progeny and contributes to developmental delay associated with failed maternal epigenetic reprogramming. RNA sequencing revealed extensive overlap between MES-4- and LSL-1-dependent transcriptional programs and demonstrated that ectopic expression of *lsl-1* itself depends on MES-4. Together, our findings support a model in which MES-4 promotes ectopic expression of LSL-1, which in turn contributes to germline-associated transcriptional programs in somatic tissues.

## MATERIALS AND METHODS

### Strains

All *Caenorhabditis elegans* strains were grown and maintained at 20°C under standard conditions, as previously described (Brenner 1974). Strains used were: N2: Bristol wild type strain provided by the *Caenorhabditis* Genetics Center; the *C. elegans spr-5 (by101)(I)* strain was provided by R. Baumeister (Albert Ludwig University of Freiburg, Germany); MT13293: *met-2 (n4256)(III)* strain was provided by R. Horvitz (Massachusetts Institute of Technology, MA, USA); the *spr-5 (by101)(I)*/*tmC27[unc-75(tmls1239)](I); met-2 (n4256) (III*)/qC1 [qls26 (lag2::gfp+ rol-6(su1006))](III) strain was created to maintain *spr-5 (by101)( I); met-2 (n4256)(III)* double-mutant animals as balanced heterozygotes (Carpenter et al. 2021); The *lsl-1(*syb3772*(lsl-1::gfp::ha::6xhis))(V)* strain was provided by C. Wicky (Rodriguez-Crespo et al. 2022) and crossed to the *met-2 (n4256) (III*)/qC1 [qls26 (lag2::gfp+ rol-6(su1006))](III) strain to maintain *met-2 (n4256) (III*)/qC1 [qls26 (lag2::gfp+ rol-6(su1006))](III); *lsl-1(*syb3772*(lsl-1::gfp::ha::6xhis))(V)* mutant animals as balanced heterozygotes.

### Scoring developmental delay

*C. elegans* adult hermaphrodites were allowed to lay embryos for 2-4 hours and then removed to synchronize the development of progeny. Progeny were then imaged and scored for development to the adult stage at 72 hours after the synchronized lay.

### RNA sequencing and analysis

Total RNA was isolated using TRIzol reagent (Invitrogen) from 1000-2000 starved L1 larvae born at 20°C overnight on unseeded NGM plates. Total RNA was sent to Georgia Genomics and Bioinformatics Core (University of Georgia, Athens, Georgia) for standard Poly-A RNA-seq services (Illumina Nextseq, 50bp paired-end reads). Raw sequencing reads were checked for quality using FastQC (Wingett and Andrews 2018), filtered using Trimmomatic (Bolger et al. 2014), and remapped to the *C. elegans* transcriptome (ce11, WBcel235) using HISAT2 (Kim et al. 2015). Read count by gene was obtained by FeatureCounts (Liao et al. 2014) . Differentially expressed transcripts (significance threshold, Wald test, p-adj < 0.05 with a log2 fold change >1 for up regulated gene and <-1 for down regulated) were determined using DESEQ2 (v.2.11.40.2) (Love et al. 2014). Transcripts per million (TPM) values were calculated from raw data obtained from FeatureCounts output. Subsequent downstream analysis was performed using R with normalized counts and p-values from DESEQ2 (v.2.11.40.2). Heatmaps were produced using the ComplexHeatmap R Package (Gu et al. 2016). Data was scaled and hierarchical clustering was performed using the complete linkage algorithm. In the linkage algorithm, distance was measured by calculating pairwise distance. Volcano plots were produced using the EnhancedVolcano package (v.1.20.0). Raw and processed RNAseq files have been deposited into Gene Expression Omnibus (www.ncbi.nlm.nih.gov/geo) under accession code GSE345214. Statistical significance of gene set overlaps was determined using hypergeometric tests. The total number of genes analyzed by DESeq2 (20,447) was used as the background population for all comparisons.

### Tissue enrichment analysis (TEA)

Tissue enrichment analyses were performed using the WormBase Tissue Enrichment Analysis (TEA) tool (Angeles-Albores et al. 2016; Angeles-Albores et al. 2018). Gene lists were submitted using WormBase gene identifiers and analyzed against the *C. elegans* tissue annotation database using default parameters. Enrichment scores were calculated relative to the genomic background utilized by the TEA algorithm, and statistical significance was determined using false discovery rate (FDR)-corrected p-values generated by the enrichment tool. Tissue categories with an FDR < 0.05 were considered significantly enriched. Fold enrichment values and corresponding FDR-adjusted p-values were used to identify tissues overrepresented among genes whose ectopic expression depended on LSL-1 in *spr-5; met-2* mutants.

### RNAi methods

RNAi by feeding was carried out using clones from the Ahringer library (Kamath and Ahringer 2003). Feeding experiments were performed on RNAi plates (NGM plates containing 100 ug/ml ampicillin, 0.4mM IPTG, and 12.5ug/ml tetracycline). F0 worms were placed on RNAi plates as L3 larvae and then moved to fresh RNAi plates 48hrs later where they were allowed to lay embryos for 2-4 hrs. F0 worms were then removed from plates and sacrificed or placed on unseeded RNAi plates overnight so that starved L1 progeny could be isolated for RNAseq experiments. F1 progeny were scored 72hrs after the synchronized lay for developmental progression. For each RNAi experiment, *pos-1* RNAi was used as a positive control. Each RNAi experiment reported here *pos-1* RNAi resulted in >95% embryonic lethality, indicating that RNAi plates were optimal.

### Differential interference contrast microscopy

Worms were immobilized in 0.1% levamisole and placed on a 2% agarose pad for Differential Interference Contrast (DIC) imaging at 10x magnification.

### Immunofluorescence staining

L1 larvae were permeabilized on slides using the freeze-crack method and immediately fixed with methanol/acetone as previously described (Duerr 2013). Following fixation, slides were washed once with 1x PBST (phosphate buffer saline w/ 0.1% Tween-20) then blocked for 30 minutes in Antibody Buffer (1x PBST with 0.5% BSA and 0.01% sodium azide). Primary antibody staining to detect the LSL-1::GFP was performed overnight at room temperature using a rabbit polyclonal anti-GFP antibody (cat. ab6556, Abcam) at a 1:600 dilution. After three washes with 1x PBST, Secondary antibody staining was performed for 1 hour at room temperature using an Alexa Fluor 594-conjugated goat anti-rabbit antibody (cat. A32740, Invitrogen) at a 1:500 dilution. Following incubation with secondary antibody, slides were washed twice with 1x PBST and once with 1x PBST containing 200 ng/ml DAPI. After three washes with 1x PBST, slides were mounted in Vectashield mounting medium and imaged immediately using a 40× objective on a Zeiss LSM 900 Confocal microscope imaging system. ImageJ maximum projection was used to project z-stack images to a single plane.

## RESULTS

### LSL-1 and genes regulated by LSL-1 in the germline are ectopically expressed in the soma of *spr-5; met-2* progeny

In a previous study, we found that germline genes, particularly those maintained by the H3K36 methyltransferase MES-4, are ectopically expressed in the soma of *spr-5; met-2* mutants and contribute to developmental delay (Carpenter et al. 2021). To determine whether genes regulated by the germline transcription factor LSL-1 are similarly misexpressed, we reanalyzed this previously published RNA-seq dataset from *spr-5; met-2* mutant L1 progeny (Carpenter et al. 2021) and compared the resulting set of 1,665 ectopically expressed genes with 1,100 genes reported to be downregulated in *lsl-1* mutant germlines (Rodriguez-Crespo et al. 2022). We found that 264 genes overlapped between these datasets (hypergeometric test, *p* = 5.65 × 10⁻⁶²; Figure 1A), representing a highly significant enrichment of LSL-1-regulated genes among transcripts ectopically expressed in *spr-5; met-2* mutant soma. Importantly, the same RNA-seq dataset revealed that *lsl-1* itself is ectopically expressed in *spr-5; met-2* mutants (log2 fold change = 1.3), raising the possibility that inappropriate activation of LSL-1 contributes to the transcriptional and developmental defects observed in reprogramming-defective animals.

**Figure 1.**
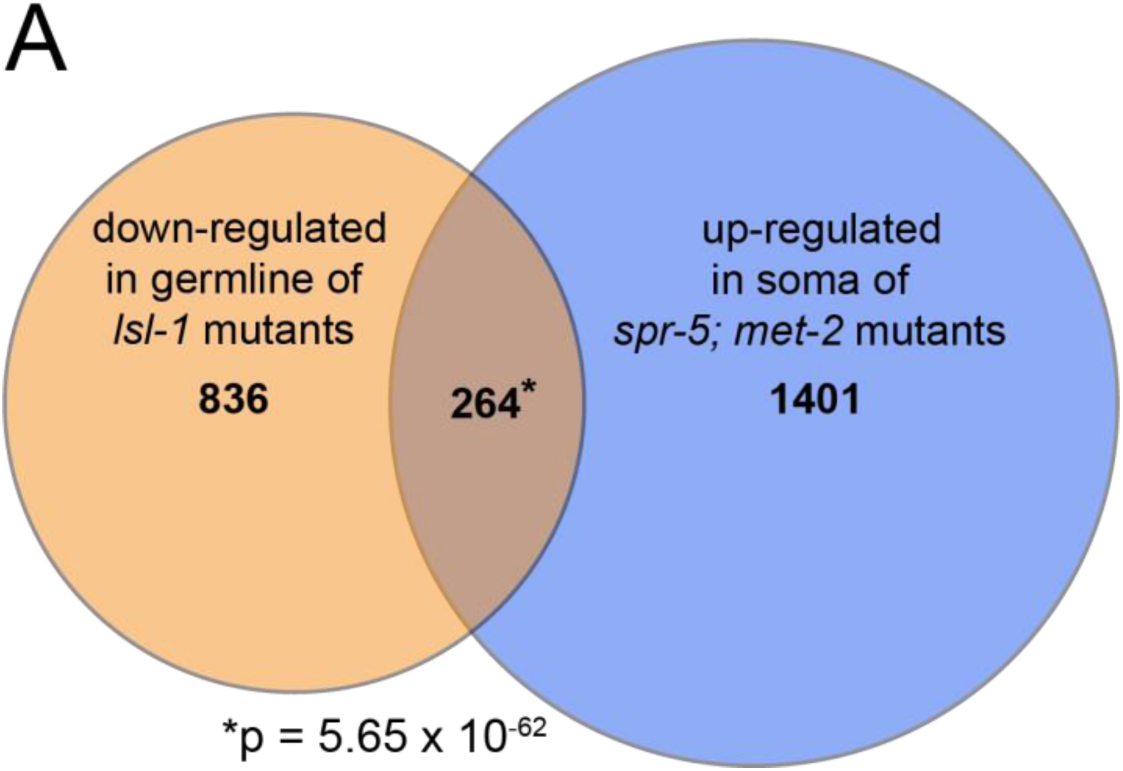
Genes ectopically expressed in *spr-5; met-2* mutant soma are enriched for genes regulated by LSL-1. (A) Overlap between genes previously identified as upregulated in *spr-5; met-2* mutant soma (1,665 genes; Carpenter et al., 2021) and genes downregulated in *lsl-1* mutant germlines (1,100 genes; Rodriguez-Crespo et al. 2022). Significant enrichment was determined by hypergeometric analysis (*p* = 5.65 × 10⁻⁶²).

Because *lsl-1* transcript levels are increased in the soma of *spr-5; met-2* mutants, we next asked whether LSL-1 protein is also ectopically expressed. To test this, we utilized an endogenously tagged *lsl-1::GFP* allele and examined L1 progeny from *met-2* mutant hermaphrodites fed either control RNAi or *spr-5* RNAi. Consistent with previous reports, LSL-1::GFP expression in control animals was restricted to the primordial germ cells Z2 and Z3 (Rodriguez-Crespo et al. 2022) ; Figure 2A-C). In contrast, progeny lacking maternal SPR-5 and MET-2 activity exhibited broad somatic expression of LSL-1::GFP throughout the body of the animal (Figure 2D-F). Together, these results demonstrate that both LSL-1 and genes normally regulated by LSL-1 in the germline are inappropriately expressed in somatic tissues in the absence of proper maternal epigenetic reprogramming.

**Figure 2.**
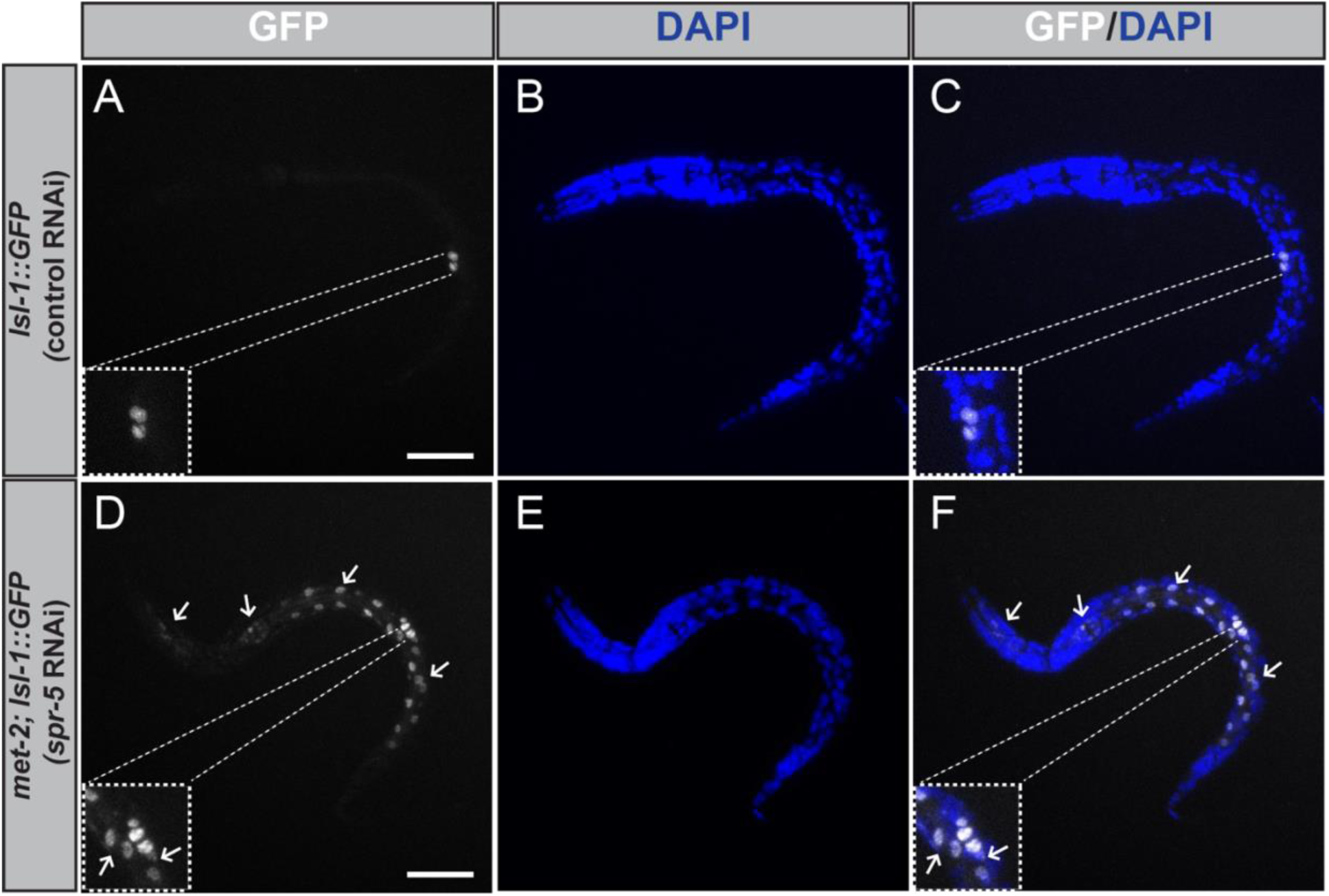
LSL-1 is ectopically expressed in the soma of *spr-5; met-2* progeny. (A-F) Representative 40x immunofluorescence confocal microscopy images of *lsl-1::GFP* in L1 larvae from hermaphrodites fed control RNAi (A-C) and *met-2; lsl-1::GFP* L1 larvae from hermaphrodites fed *spr-5* RNAi (D-F). DAPI was used to stain nuclear chromatin. Insets denote the location of the primordial germ cells, Z2 and Z3. White arrows denote somatic expression of *lsl-1::GFP*. Scale bars: 10 µm.

### Knocking down LSL-1 partially suppresses developmental delay in *spr-5; met-2* progeny

The observation that LSL-1 target genes are ectopically expressed in the soma of *spr-5; met-2* mutants led us to ask whether LSL-1 contributes to the developmental defects associated with failed maternal reprogramming. To test this, *spr-5; met-2* hermaphrodites were fed *lsl-1* RNAi and progeny were scored for developmental progression 72 hours after a synchronized lay (Figure 3). To begin with, we assessed the effectiveness of RNAi knockdown by feeding *lsl-1::GFP* animals *lsl-1* RNAi and examining immunofluorescence. Approximately 90% of animals lacked detectable GFP signal, confirming efficient depletion of LSL-1 (Supplemental Figure 1). As previously observed, nearly all wild-type progeny reached adulthood by 72 hours (Figure 3A, E), while only 9% of *spr-5; met-2* progeny fed control RNAi developed to adults (Figure 3B, E). Consistent with previous findings, knockdown of *mes-4* strongly suppressed the developmental delay phenotype, allowing 79% of progeny to reach adulthood (Carpenter et al. 2021); Figure 3C, E). Knockdown of *lsl-1* also significantly rescued developmental progression, with 52% of progeny reaching adulthood by 72 hours (Figure 3D, E).

**Figure 3.**
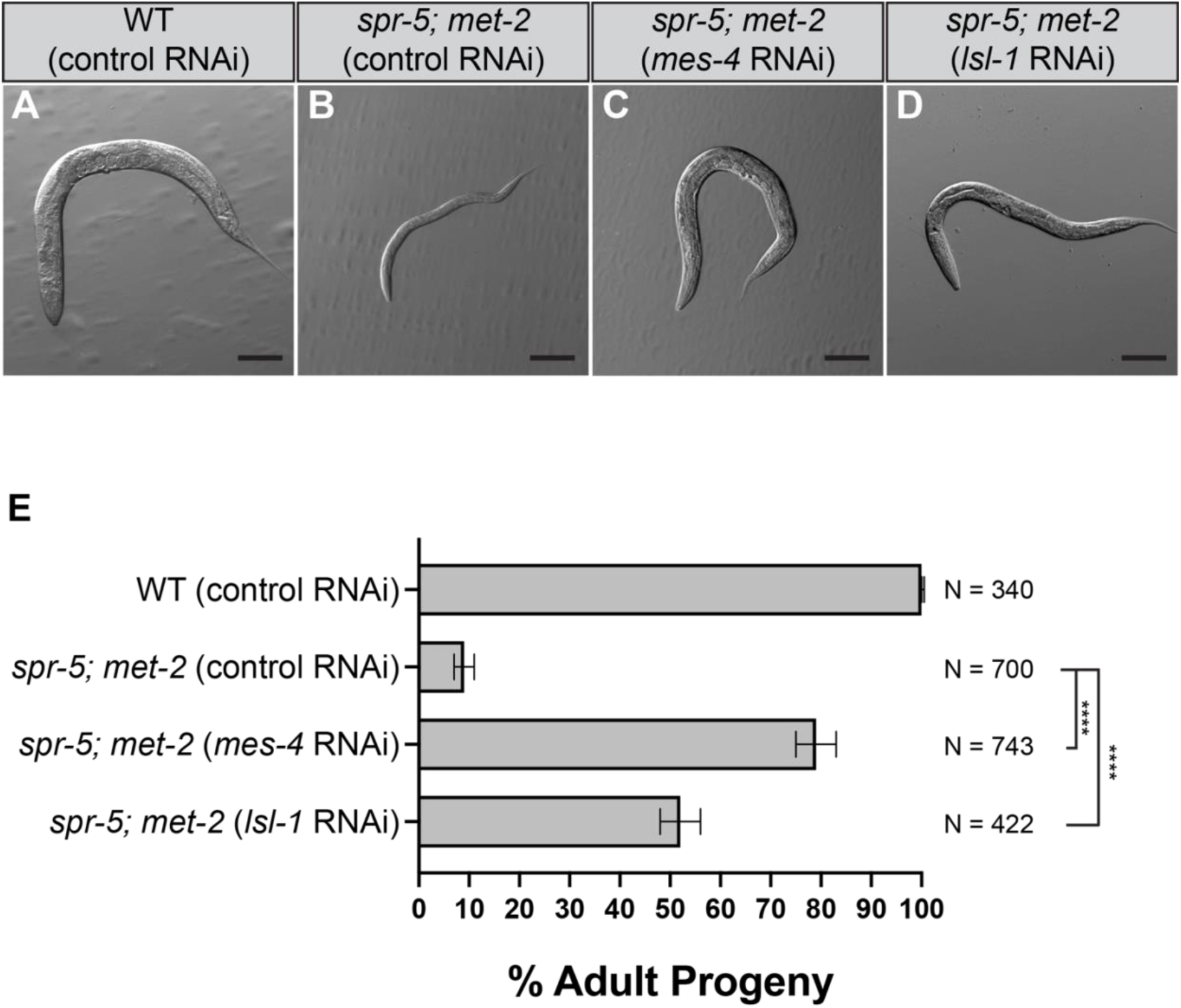
Knocking down LSL-1 partially rescues developmental delay in *spr-5; met-2* progeny. DIC images of wild type (A) or *spr-5; met-2* (B-D) progeny from hermaphrodite parents treated L4440 vector only (control) RNAi (A-B), *mes-4* RNAi (C), or *lsl-1* RNAi (D) 72 hours post synchronized lay. Scale bar: 100μm. (E) Quantification of the number of progeny (represented as % Adult Progeny) from A-D that made it to adults by 72 hours. Error bars represent the standard deviation of the mean from three biological replicates. N represents the total number of progeny from 30-40 hermaphrodites scored across independent experiments. (unpaired t-test, **** represent a p-value <0.0001).

### LSL-1 contributes to ectopic expression of a MES-4-dependent germline-associated transcriptional program in *spr-5; met-2* mutant soma

To determine the extent to which ectopic gene expression in *spr-5; met-2* mutants depends on MES-4 and LSL-1, we performed RNA sequencing on starved L1 progeny from *spr-5; met-2* animals fed control RNAi, *mes-4* RNAi, or *lsl-1* RNAi (Figure 4, Supplemental Figure 2). Because starved L1 larvae contain only two germ cells that have not yet activated germline transcription, this developmental stage allows analysis of transcriptional defects occurring in somatic tissues. Among the 1,323 genes upregulated in the soma of *spr-5; met-2* mutants, 1,012 exhibited reduced expression following *mes-4* RNAi and 818 exhibited reduced expression following *lsl-1* RNAi (Figure 4A). Comparison of these datasets identified 657 overlapping genes, representing a highly significant overlap between MES-4- and LSL-1-dependent transcriptional programs (hypergeometric test, *p* < 4.1 × 10⁻^803^; Figure 4A). These findings indicate that most genes dependent on LSL-1 are also dependent on MES-4. Furthermore, *lsl-1* itself was among the MES-4-dependent genes. While *lsl-1* expression was elevated in *spr-5; met-2* mutants (log2 fold change = 1.3), expression was reduced to near wild-type levels following *mes-4* RNAi treatment (log2 fold change = -0.17; Figure 4D), suggesting that ectopic expression of *lsl-1* depends on MES-4.

**Figure 4.**
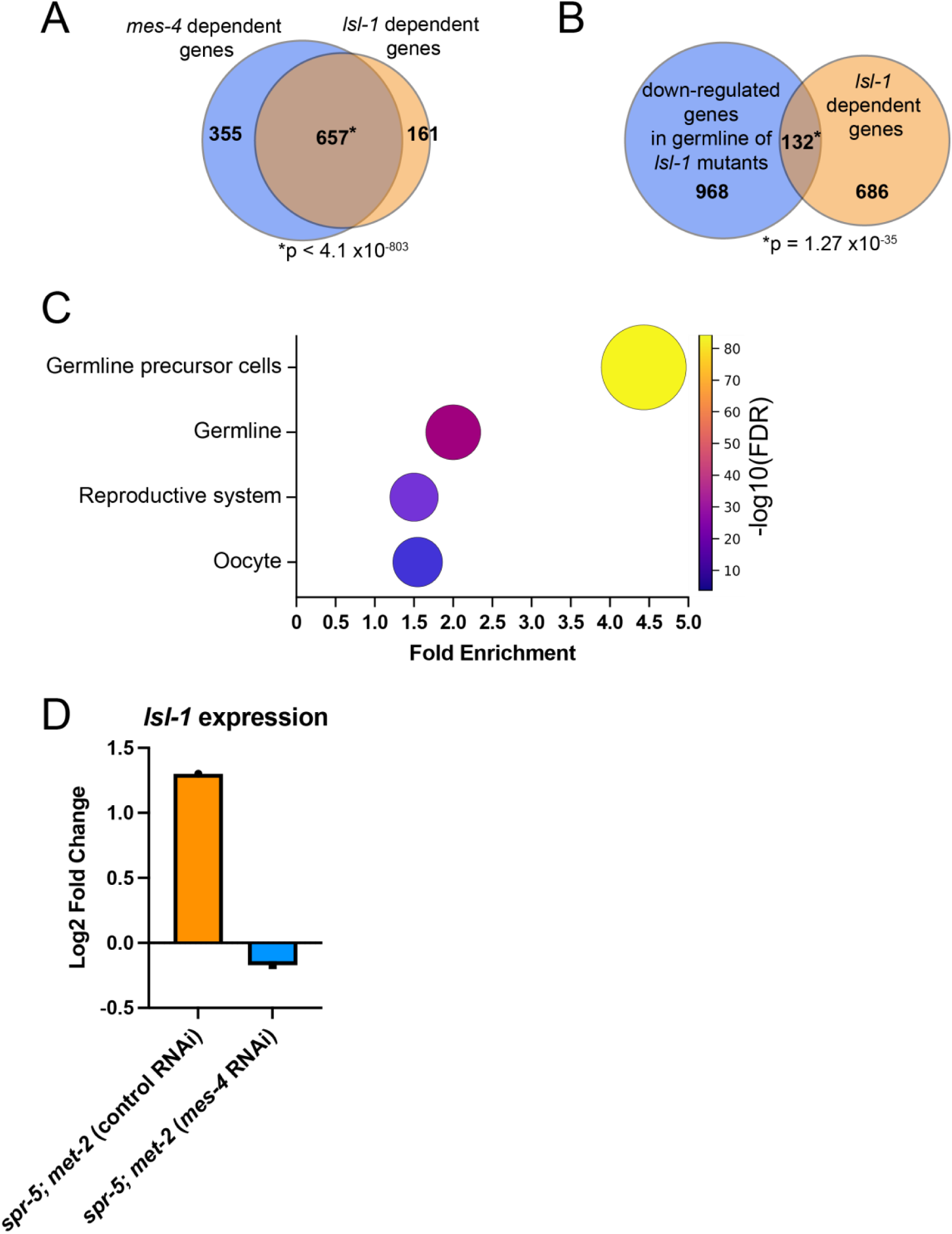
MES-4 and LSL-1 promote a shared germline-associated transcriptional program in *spr-5; met-2* mutant soma. (A) Overlap between genes whose ectopic expression in *spr-5; met-2* mutants was reduced by *mes-4* RNAi (1,012 genes) and genes whose ectopic expression was reduced by *lsl-1* RNAi (818 genes). A total of 657 genes overlapped between these datasets (hypergeometric test, *p* < 4.1 × 10⁻^803^), indicating extensive overlap between MES-4- and LSL-1-dependent transcriptional programs. (B) Overlap between genes whose expression was reduced by *lsl-1* RNAi in *spr-5; met-2* mutants (818 genes) and genes previously identified as downregulated in the germline of *lsl-1* mutants (1,100 genes). A total of 132 genes overlapped between these datasets (hypergeometric test, *p* = 1.27 × 10⁻^35^), demonstrating significant enrichment of bona fide LSL-1-regulated genes among the LSL-1-dependent transcriptional program. (C) Tissue Enrichment Analysis (TEA) of genes whose ectopic expression depended on LSL-1. The x-axis indicates fold enrichment relative to genomic expectation. Bubble size is proportional to fold enrichment and bubble color denotes statistical significance (-log10 FDR). Genes dependent on LSL-1 were significantly enriched for expression in germline precursor cells, the germ line, reproductive tissues, and oocytes. (D) Expression of *lsl-1* in *spr-5; met-2* mutants following control or *mes-4* RNAi treatment. RNA-seq analysis revealed that ectopic *lsl-1* expression observed in *spr-5; met-2* mutants (log2 fold change = 1.3) was reduced to near wild-type levels following *mes-4* RNAi (log2 fold change = -0.17), indicating that ectopic expression of *lsl-1* depends on MES-4.

To determine whether genes responsive to *lsl-1* RNAi correspond to genes normally regulated by LSL-1 in the germline, we compared the 818 LSL-1-dependent genes to published transcriptomic datasets from *lsl-1* mutants. We identified 132 genes shared between the LSL-1-dependent gene set and 1,100 genes previously reported to be downregulated in *lsl-1* mutant germlines (hypergeometric test, *p* = 1.27 × 10⁻^35^; Rodriguez-Crespo et al. 2022; Figure 4B). This significant overlap indicates that a subset of genes ectopically expressed in *spr-5; met-2* mutant soma corresponds to genes that normally require LSL-1 for expression in the germline.

We next investigated whether genes dependent on LSL-1 were associated with specific tissue-expression programs. Tissue Enrichment Analysis (TEA) revealed significant enrichment for genes expressed in germline precursor cells, the germ line, reproductive tissues, and oocytes (Figure 4C). Germline precursor cells represented the most strongly enriched category (4.4-fold enrichment, FDR = 4.8 × 10⁻⁸⁵), followed by germ line (2.0-fold enrichment, FDR = 3.7 × 10⁻³⁶), reproductive system (1.5-fold enrichment, FDR = 2.1 × 10⁻¹⁷), and oocyte (1.6-fold enrichment, FDR = 1.9 × 10⁻⁴). These findings indicate that genes dependent on ectopic LSL-1 are strongly enriched for germline-associated expression programs. Consistent with the possibility that some of these transcriptional effects are direct, 52 genes overlapped between the 818 LSL-1-dependent genes and a published set of 296 genes that are both promoter-bound by LSL-1 and downregulated in *lsl-1* mutant germlines (hypergeometric test, *p* =1.52× 10⁻^19^; Rodriguez-Crespo et al. 2022; Supplemental Figure 3). Together, these findings suggest that both direct and indirect mechanisms contribute to the LSL-1-dependent transcriptional program observed in *spr-5; met-2* mutants and support a model in which MES-4 promotes ectopic expression of LSL-1, resulting in a shared germline-associated transcriptional program following failed maternal epigenetic reprogramming (Figure 5).

**Figure 5.**
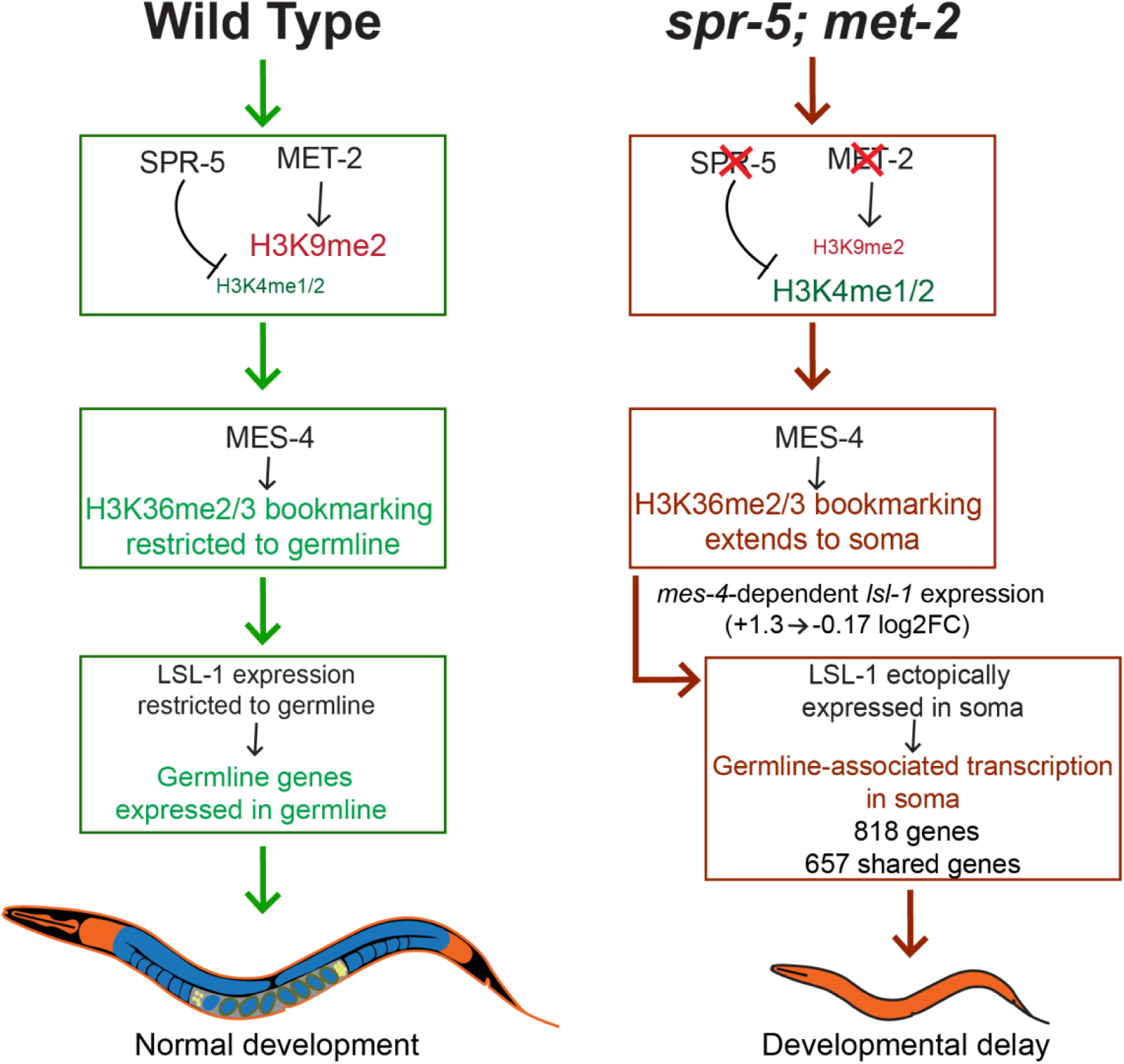
Proposed model for how failed maternal epigenetic reprogramming leads to ectopic germline-associated transcription and developmental delay. Maternal SPR-5 and MET-2 normally reprogram inherited chromatin states by removing H3K4me1/2 and establishing H3K9me2 during embryogenesis. In wild-type animals, MES-4-dependent H3K36me2/3 bookmarking and LSL-1 expression are restricted to germ cells, resulting in germline gene expression exclusively within the germline and normal development. In *spr-5; met-2* mutants, persistent H3K4me1/2 and loss of proper H3K9me2 establishment allow MES-4-dependent H3K36me2/3 bookmarking to extend into somatic tissues. Ectopic MES-4 activity promotes LSL-1 expression and germline-associated transcription in the soma. The extensive overlap between MES-4- and LSL-1-dependent gene sets supports a model in which these factors cooperate to promote ectopic germline transcription, contributing to developmental delay.

## DISCUSSION

Proper establishment and maintenance of cell identity requires coordinated interactions between chromatin regulators and sequence-specific transcription factors. In *C. elegans*, maternal reprogramming factors reset inherited chromatin states following fertilization and prevent inappropriate activation of germline-specific transcriptional programs in somatic tissues. Previous studies demonstrated that loss of the maternal reprogramming factors SPR-5 and MET-2 results in ectopic germline gene expression and severe developmental delay, and that these defects depend in part on the chromatin regulator MES-4 (Katz et al. 2009; Greer et al. 2014; Kerr et al. 2014; Carpenter et al. 2021). However, whether germline-specific transcription factors contribute to the ectopic transcriptional program arising from failed maternal epigenetic reprogramming remained unclear. Here, we demonstrate that the germline transcription factor LSL-1 becomes ectopically expressed in somatic tissues of *spr-5; met-2* progeny and contributes to both developmental delay and ectopic germline-associated transcription.

Our findings suggest that failed maternal reprogramming disrupts germline-soma boundaries at multiple levels of gene regulation. Consistent with previous work, knockdown of *mes-4* reduced expression of a large fraction of genes ectopically expressed in *spr-5; met-2* mutants and strongly rescued developmental progression (Carpenter et al. 2021). Knockdown of *lsl-1* similarly rescued developmental delay and reduced expression of 818 ectopically expressed genes. Notably, 657 genes were shared between the MES-4- and LSL-1-dependent gene sets, indicating extensive overlap between the transcriptional programs regulated by these factors. Furthermore, *lsl-1* itself was among the MES-4-dependent genes. Ectopic expression of *lsl-1* observed in *spr-5; met-2* mutants was reduced to near wild-type levels following *mes-4* RNAi treatment, suggesting that MES-4 contributes to ectopic germline transcription at least in part through activation of LSL-1 (Figures 4 and 5).

Several independent lines of evidence indicate that genes dependent on LSL-1 are enriched for germline-associated functions. First, genes previously shown to require LSL-1 in the germline were significantly enriched among genes whose ectopic expression depended on LSL-1 in *spr-5; met-2* mutants. Second, Tissue Enrichment Analysis revealed strong enrichment for germline precursor cell, germline, reproductive system, and oocyte expression programs. Germline precursor cells were particularly enriched, suggesting that the genes most dependent on ectopic LSL-1 correspond to core germline transcriptional programs. Together, these findings indicate that ectopic activation of LSL-1 contributes specifically to germline-associated transcription rather than broadly altering somatic gene expression.

Although many genes depended on LSL-1 for ectopic expression in *spr-5; met-2* mutants, only a subset corresponded to previously identified direct LSL-1 targets. Nevertheless, direct LSL-1 targets were significantly enriched among the LSL-1-dependent gene set, supporting the conclusion that at least part of the ectopic transcriptional program is regulated directly by LSL-1. One possibility is that a relatively small number of direct targets function as upstream regulators that subsequently influence broader transcriptional networks. Alternatively, ectopic LSL-1 may only activate a subset of its normal target genes because the chromatin environment required for full LSL-1 activity is present in germ cells but absent from somatic tissues (Rodriguez-Crespo et al. 2022). In this model, MES-4-dependent chromatin states may create a permissive environment that enables ectopic expression of LSL-1 and a subset of its targets, while additional germline-specific regulatory factors limit activation of the complete germline transcriptional program.

An important implication of this work is that inherited chromatin states alone may be insufficient to explain the developmental consequences of failed maternal epigenetic reprogramming. Much attention has focused on ectopic histone modifications and chromatin memory as drivers of inappropriate germline gene expression outside of the germline (Furuhashi et al. 2010; Rechtsteiner et al. 2010; Carpenter et al. 2021). Our findings suggest that lineage-restricted transcription factors represent an additional layer of regulation that reinforces and amplifies aberrant transcriptional states once they become established. As summarized in Figure 5, we propose that loss of SPR-5 and MET-2 permits MES-4-dependent activation of germline-associated chromatin states, leading to ectopic expression of LSL-1 and activation of a shared germline-associated transcriptional program that contributes to developmental delay.

In summary, we show that the germline transcription factor LSL-1 is ectopically expressed in the soma of *spr-5; met-2* mutants and contributes to both developmental delay and ectopic germline-associated transcription. Our results identify LSL-1 as a transcriptional effector acting downstream of MES-4 and reveal how chromatin regulators and lineage-restricted transcription factors cooperate to disrupt cell fate boundaries following failed maternal epigenetic reprogramming. These findings reveal how inherited chromatin states and lineage-restricted transcription factors cooperate to disrupt cell fate boundaries following failed maternal epigenetic reprogramming.

---

**Supplemental Figure 1.**
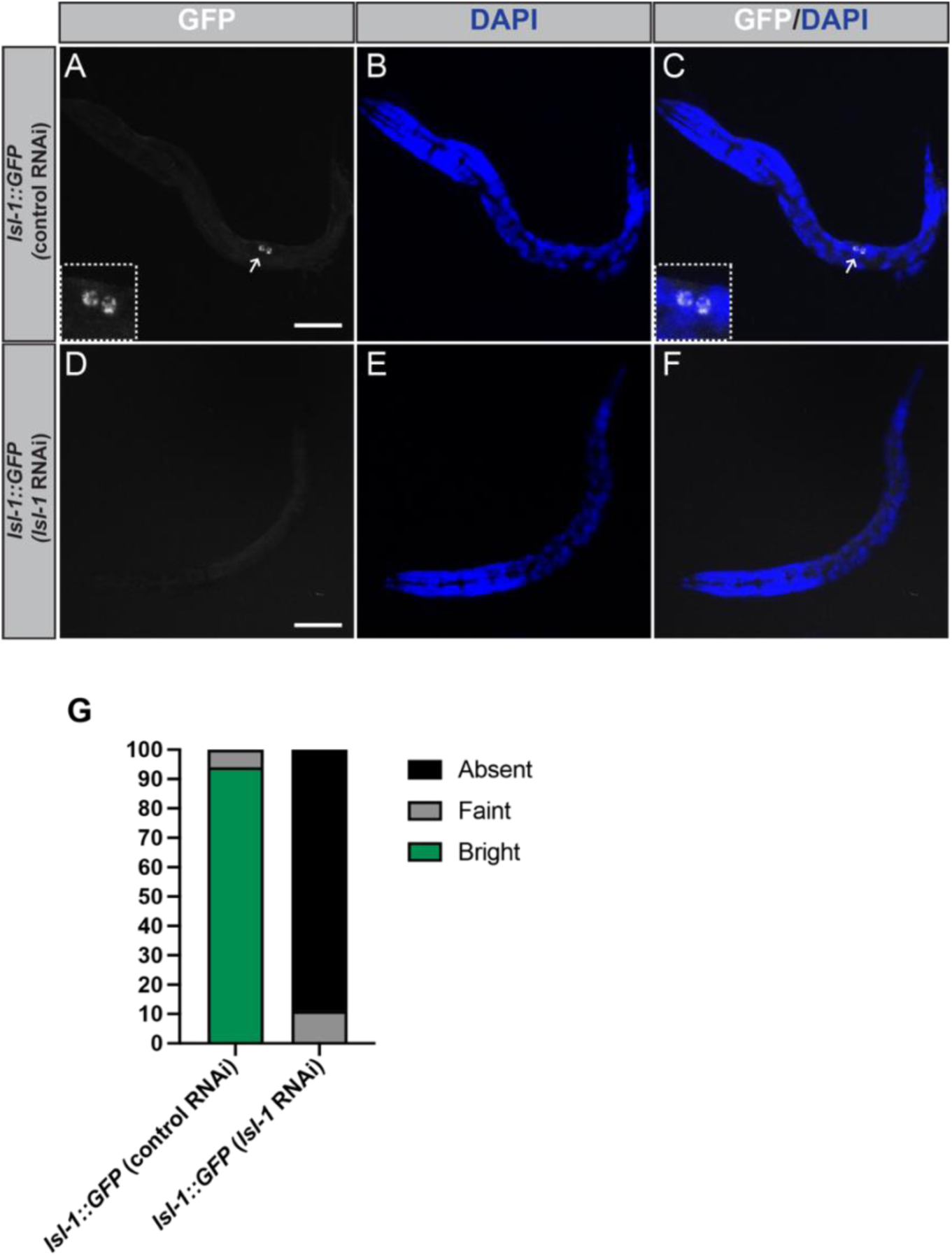
*lsl-1* RNAi efficiently depletes LSL-1::GFP protein expression. Representative 40× immunofluorescence confocal microscopy images of *lsl-1::GFP* L1 larvae from hermaphrodites fed control RNAi (A-C) or *lsl-1* RNAi (D-F). GFP signal was detected using an anti-GFP antibody and nuclei were visualized with DAPI. In control animals, LSL-1::GFP expression was restricted to the primordial germ cells Z2 and Z3 (A-C, inset and arrow), consistent with previously reported germline-specific expression. In contrast, L1 larvae exposed to *lsl-1* RNAi exhibited little or no detectable GFP signal (D-F), indicating efficient depletion of LSL-1 protein. Scale bars: 10 μm. (G) Quantification of animals exhibiting detectable LSL-1::GFP signal following control or *lsl-1* RNAi treatment. Approximately 90% of animals exposed to *lsl-1* RNAi lacked detectable GFP expression, confirming effective knockdown of LSL-1.

**Supplemental Figure 2.**
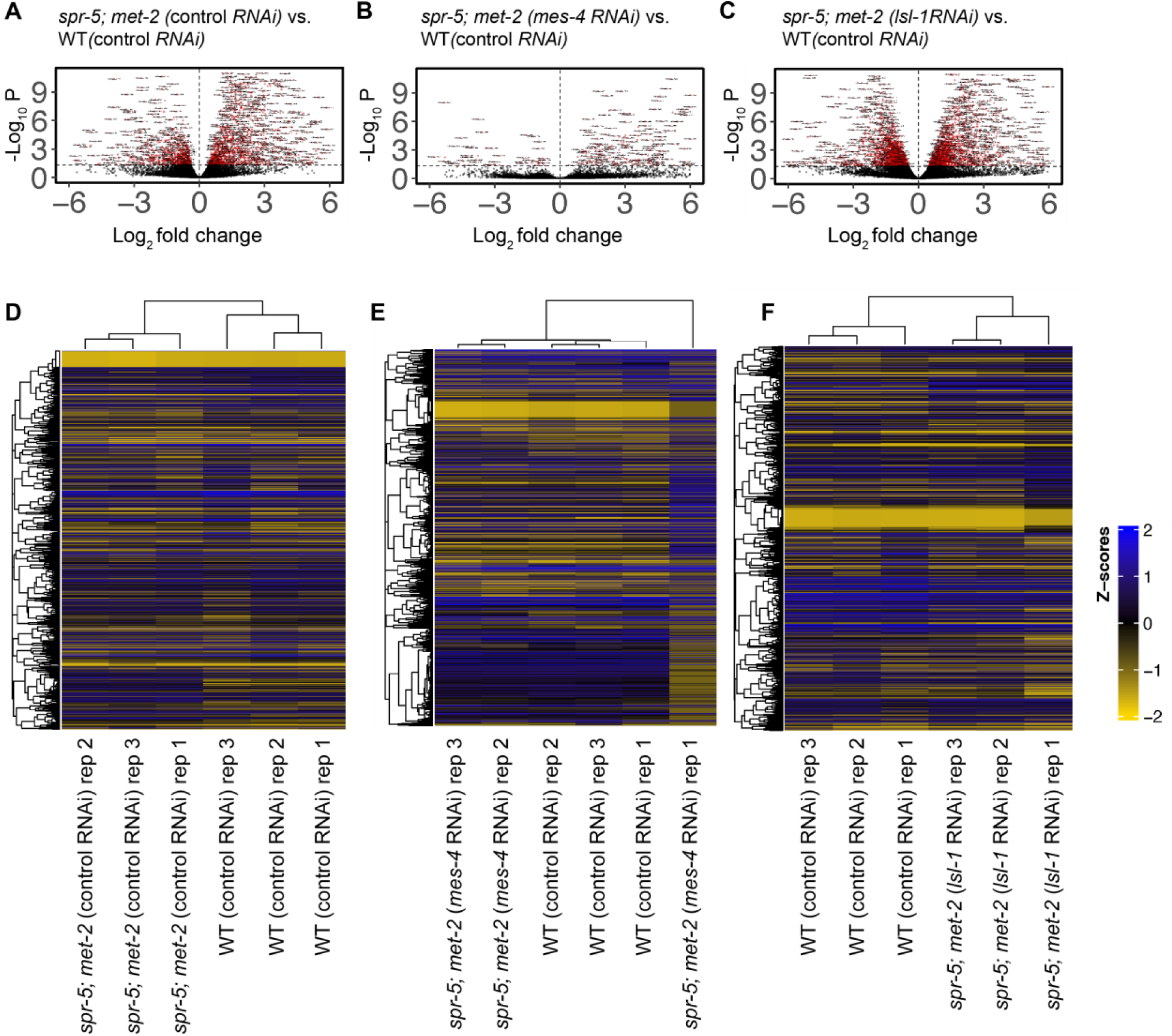
Differential expression and replicate comparison of RNAseq experiments performed on wild type, *spr-5; met-2* progeny fed either control, *mes-4,* or *lsl-1* RNAi. Volcano plot of log2 fold changes in gene expression (x-axis) by statistical significance (-Log_10_ P-value; y-axis) in L1 progeny of *spr-5; met-2* hermaphrodites fed L4440 (control) RNAi (A), *spr-5; met-2* hermaphrodites fed *mes-4* RNAi (B), and *spr-5; met-2* hermaphrodites fed *lsl-1* RNAi (C) compared to L1 progeny wild type (WT) hermaphrodites fed control RNAi. Heatmap of differentially expressed RNA-seq transcripts between L1 progeny of *spr-5; met-2* hermaphrodites fed control RNAi (D), *spr-5; met-2* hermaphrodites fed *mes-4* RNAi (E), and *spr-5; met-2* hermaphrodites fed *lsl-1* RNAi (F). Data was scaled and hierarchical clustering was performed using complete linkage algorithm, with distance measured by calculating pairwise distance. Higher (blue) and lower (yellow) expression is reported as a z-score.

**Supplemental Figure 3.**
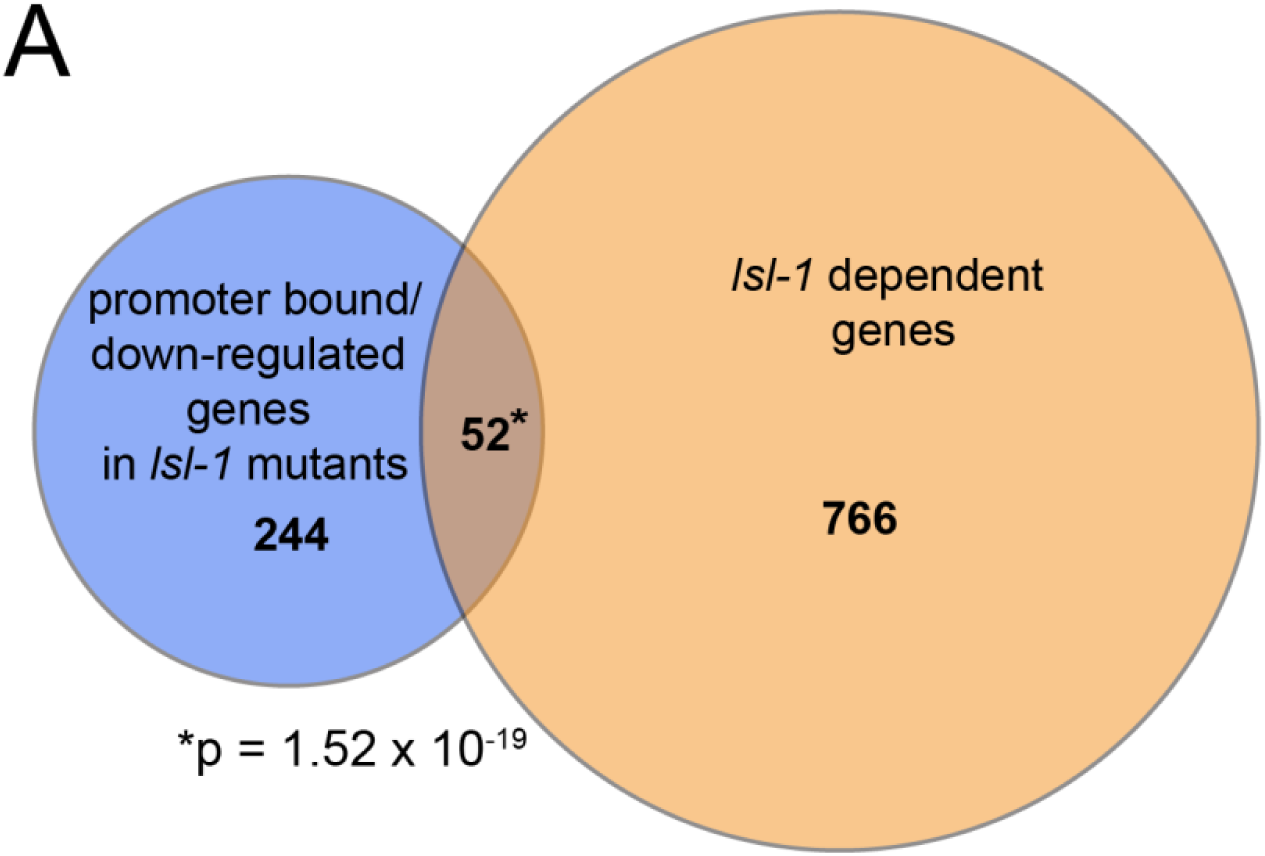
A subset of LSL-1-dependent genes correspond to previously identified direct LSL-1 targets. (A) Overlap between genes whose ectopic expression was reduced by *lsl-1* RNAi in *spr-5; met-2* mutants (818 genes) and a published set of genes that are both promoter-bound by LSL-1 and downregulated in *lsl-1* mutant germlines (296 genes). Fifty-two genes overlapped between these datasets (hypergeometric test, *p* = 1.52 × 10^⁻19^), representing a significant enrichment of direct LSL-1 targets among the LSL-1-dependent transcriptional program. These genes represent high-confidence candidate direct targets of ectopic LSL-1 activity in *spr-5; met-2* mutant soma.

## Supplemental Files

**Supplemental File S1. Genes ectopically expressed in *spr-5; met-2* mutant soma.** List of differentially expressed genes identified by RNA-seq analysis of starved L1 progeny from *spr-5; met-2* mutants compared with wild-type controls. Included are WormBase gene identifiers, gene names, log2 fold changes, and adjusted P-values. Differential expression was defined as adjusted P < 0.05 and absolute log2 fold change > 1.

**Supplemental File S2. MES-4-dependent genes ectopically expressed in spr-5; met-2 mutant soma.** Genes whose ectopic expression in *spr-5; met-2* mutant progeny was reduced following *mes-4* RNAi treatment. Included are WormBase gene identifiers, gene names, and differential expression statistics from RNA-seq analyses of wild-type, *spr-5; met-2*, and *spr-5; met-2*; *mes-4* RNAi progeny. Genes were classified as MES-4-dependent based on reduced expression following *mes-4* RNAi relative to the ectopic expression observed in *spr-5; met-2* mutants.

**Supplemental File S3. LSL-1-dependent genes ectopically expressed in spr-5; met-2 mutant soma.** Genes whose ectopic expression in *spr-5; met-2* mutant progeny was reduced following *lsl-1* RNAi treatment. Included are WormBase gene identifiers, gene names, and differential expression statistics from RNA-seq analyses of wild-type, *spr-5; met-2*, and *spr-5; met-2*; *lsl-1* RNAi progeny. Genes were classified as LSL-1-dependent based on reduced expression following *lsl-1* RNAi relative to the ectopic expression observed in *spr-5; met-2* mutants.

**Supplemental File S4. Candidate direct LSL-1 targets ectopically expressed in *spr-5; met-2* mutant soma.** Genes with promoter-proximal LSL-1 binding and reduced expression in *lsl-1* mutants were obtained from Rodriguez et al. (2022). Candidate direct LSL-1 targets were defined as genes from this set whose ectopic expression in *spr-5; met-2* mutant progeny was reduced following *lsl-1* RNAi treatment.

## Acknowledgements

We thank Chantal Wicky (University of Fribourg) for generously providing the *lsl-1(syb3772[lsl-1::gfp::ha::6xhis])* strain used in this work. Some strains were provided by the *Caenorhabditis* Genetics Center (CGC), which is funded by the NIH Office of Research Infrastructure Programs (P40 OD010440). We also thank members of the Carpenter laboratory and David Katz (Emory University) for helpful discussions and feedback on the project. We also acknowledge WormBase for providing genomic and gene annotation resources that facilitated data analysis and interpretation.

## Data Availability Statement

Strains and reagents used in this study are available upon request. Raw and processed RNAseq files from experiments performed in this study have been deposited into Gene Expression Omnibus (www.ncbi.nlm.nih.gov/geo) under accession code GSE345214. All previously published datasets analyzed in this study are publicly available, including GEO accession GSE154649 (Carpenter et al., 2021), ArrayExpress accession E-MTAB-11199 (Rodríguez-Crespo et al., 2022 RNA-seq), and LSL-1 ChIP-seq datasets available through the modERN project repository.

